# ALE Meta-Analysis Reveals Neural Substrates for the Impact of Prematurity on Executive Functioning in Children and Adults

**DOI:** 10.1101/2025.06.24.661281

**Authors:** Anna E. Youngkin, Gabriella Snetkov, Cara Grady, Meghan H. Puglia, Katheryn Frazier, Kevin A. Pelphrey, Karen Fairchild, Howard P. Goodkin, Tanya M. Evans

## Abstract

Premature birth has known impacts on brain development, leading to sustained differences in cognitive function throughout the lifespan. Despite known deficits in executive functioning (EF) within individuals born premature, the extent to which neural engagement during executive functioning tasks differs between those born preterm and full-term is not fully understood. Additionally, it is unknown whether regions of differential engagement are the same in children and adults. This meta-analysis synthesizes fMRI results of activation differences between preterm and full-term subjects during executive functioning tasks in adult and child groups separately. Our results indicate that differences in neural engagement during EF tasks differ between pre-term (PT) and full term (FT) individuals in both age groups. Moreover, the regions affected contribute to well-known brain networks, including the fronto-striatal circuitry, the default mode network (DMN), and the salience network, all of which subserve broad EF capabilities. We found no differences between child and adult maps in a direct contrast, suggesting that effects of prematurity on executive functioning may persist from childhood into adulthood, although these findings should be interpreted in context of methodological limitations and potential confounding factors. This meta-analysis provides greater insight into the neural mechanisms behind EF disruption following premature birth.

**Highlights:**

- Differences in neural activation during executive function tasks exist in both children and adults with a history of premature birth.
- PT children show hyperactivity in fronto-striatal regions while PT adults show differential engagement of default mode network regions.

## Introduction

The United States has the highest incidence of preterm birth–defined as birth prior to 37 weeks gestational age (GA)–relative to countries of similar economic status, with 1 in 10 babies born prematurely (Purisch & Gyamfi-Bannerman, 2017). Despite the survival of extremely preterm infants increasing substantially over the last decade (Stoll et al., 2015), advancements in prenatal and neonatal care have not led to demonstrated improved cognitive and academic outcomes (Cheong et al., 2020). As a result, children born premature are less likely to be ready for school entry (Taylor et al., 2022) and have worse academic outcomes across curriculum areas compared to term-born peers (Allotey et al., 2018; Pritchard et al., 2009). Deficits in executive functioning (EF) are particularly relevant following PT birth as they are shown to persist across the lifespan (Blasco et al., 2024; Burnett et al., 2013; Schnider et al., 2020; Sølsnes et al., 2014). Importantly, EF deficits in PT adults are associated with decreased years of education, lower socio-economic status, and higher rates of unemployment (Kim et al., 2022; Kroll et al., 2017). Therefore, understanding the mechanisms leading to changes in EF following PT birth has important implications for academic outcomes and quality of life.

Executive functioning is a broad term encompassing multiple cognitive processes that aid in planning and managing appropriate behavior, often broken down into inhibition, working memory, and cognitive flexibility (Diamond, 2013). The subdomains of EF have distinct developmental trajectories, with inhibition reaching adult levels earliest in childhood (Huizinga et al., 2006; Johnstone et al., 2007) followed by cognitive flexibility, while working memory skills continue to improve into early adulthood (McAuley & White, 2011). While these three main subdomains of EF are often considered distinct and can be isolated from each other in behavioral tasks (Fisk & Sharp, 2004; Miyake et al., 2000), their development is often viewed as interconnected and interdependent (Brocki & Bohlin, 2004; Buttelmann & Karbach, 2017).

Children born preterm exhibit persistent deficits within these domains of EF. They have been shown to have lower cognitive, motor, and academic performance scores, while exhibiting higher scores on behavior assessments indicative of problems (Allotey et al., 2017). When EF component scores of the BSID-III assessment were compared between term and preterm, preterm children were found to have statistically lower scores in categories of attention, working memory, and plan/organize (Blasco et al., 2024). Prematurely born adults performed worse than controls on neurocognitive tests measuring EF, such as HSCT, COWAT, IED, and TMT-B (Kroll et al., 2017). Additionally, very preterm-born children have been shown to recruit additional prefrontal areas compared to controls during a spatial working memory task (Pia-Maria et al., 2017). Enhanced deactivation of the default mode network was observed in very preterm-born adults during the difficult 2-back condition of a verbal N-back task, demonstrating the possibility of load-dependent differences in brain activity (Daaman et al., 2014). These findings underscore that EF impairments following preterm birth are both behavioral and neural in nature, suggesting that PT individuals may engage EF networks differently across development.

Moreover, the subdivision of EF skills becomes less clearly defined behaviorally as EF skills mature: throughout development, the tasks needed to assess EF grow more complex. As a result, tasks utilized to measure a single domain of EF often begin to inadvertently assess multiple converging subdomains at once (Asato et al., 2006). Additionally, the neural substrates of EF are distributed among multiple distinct yet interconnected neural circuits (Witt et al., 2021). While individual subdomains do rely upon unique neural signatures (Lemire-Rodger et al., 2019), there are shared neuronal networks across EF subdomains (Saylik et al., 2022; Sylvester et al., 2003), which may be thought of as a common EF shared factor (Friedman & Miyake, 2017). Importantly, the impact of PT birth on EF is simultaneously broad and specific, as children born PT show deficits in individual subdomains as well as within broader EF skills (Stålnacke et al., 2019). As a result, the neural basis for EF deficits following PT birth may either be within subdomains individually or due to changes in networks underlying broader EF skills.

Prior research has shown that individuals born preterm often experience persistent EF difficulties and altered neural activation during EF tasks. However, much of this work has focused on specific EF subdomains using single-task fMRI studies, making it difficult to determine whether differences reflect narrow impairments or broader disruptions to domain-general EF networks. In addition, findings across studies are inconsistent, likely due to differences in tasks, age groups, and analysis methods. Few studies directly compare PT and FT individuals across developmental stages, leaving it unclear whether neural differences remain stable or shift over time. This meta-analysis aims to address two main questions: (1) Does preterm birth affect brain activation in regions supporting common EF across tasks? and (2) Do these effects differ between childhood and adulthood? By synthesizing data across studies, we aim to clarify how PT birth impacts the neural basis of EF across development.

To address these gaps, the current meta-analysis was performed to test the hypothesis that PT birth leads to differential development of the neural substrates underlying ‘common’ EF skills, leading to broad behavioral effects across EF subdomains following preterm birth. Much of the work assessing changes to neural substrates of EF following PT birth utilize single EF tasks, leading to findings that are focused on individual subdomains. By statistically combining findings across multiple task modalities, our meta analysis aims to identify differences in broader EF networks between PT and full-term (FT) subjects. Moreover, we utilize findings from fMRI studies completed in both children (6-14yo) and adults (20-35yo). Therefore, we are able to assess whether the differences in PT and FT subjects vary between childhood and adulthood. We expect to find patterns of different neural activation in PT subjects in regions most often associated with broader EF skills in both children and adults, such as the medial prefrontal gyrus, left prefrontal cortex, and bilateral intraparietal sulci (McKenna et al., 2017; Reineberg et al., 2022; Santarnecchi et al., 2021). Results indicating significant differences in regions known to subserve specific subdomains–such as the unique role of the caudate in response inhibition (Wager et al., 2005; R. Zhang et al., 2017)–would indicate domain-specific neural deficits. The current meta analysis provides meaningful insight into the impact of PT birth on networks subserving both common EF skills and EF subdomains and examines whether the impact of PT birth differs in childhood versus adulthood.

## Methods

### Study Selection

PubMed, Embase, and Web of Science databases were searched to identify articles using the search terms ‘preterm’ or ‘premature,’ ‘fMRI,’ or ‘PET’, ‘executive function’ or ‘executive functioning,’ ‘working memory.’ ‘inhibition’ and ‘switching’. Working memory (WM) was used as an additional search term due to evidence that it may be the shared common EF factor (Lemire-Rodger et al., 2019), and that many WM papers did not include mention of executive function. Reference lists and studies citing these articles were examined using citation chaining to maximize the inclusion of all relevant studies. The final search was completed May 3, 2025. Articles were screened and assessed for eligibility according to PRISMA guidelines (Page et al., 2021). 348 total studies were imported for screening, of which 71 were duplicates. Two reviewers (AY, GS) independently completed abstract screening, and studies were deemed irrelevant if the abstract did not discuss premature birth, did not complete neuroimaging, or was focused on another biological system (e.g., cardiac development). If either reviewer deemed the study relevant, the study was moved into full-text review. 219 studies were deemed irrelevant during the abstract review, leaving 58 studies that were assessed for eligibility. The following criteria were used to determine study eligibility: (1) the study’s participants included a PT group born <37 weeks GA and a FT control group; (2) participants underwent fMRI or PET imaging during a behavioral task assessing at least one of the three EF domains; (3) coordinates of significantly activated/deactivated voxels were reported in either MNI or Talairach coordinates; (4) the study undertook and reported results from a whole-brain (as opposed to region of interest) analysis. 19 studies were ultimately included in our meta-analysis (see Table 1). The interrater reliability score was 94.3% for studies moved from the initial abstract review to full text review, and 94.1% for studies moved from the full text review to be included in the final analysis.

**Table 1:**
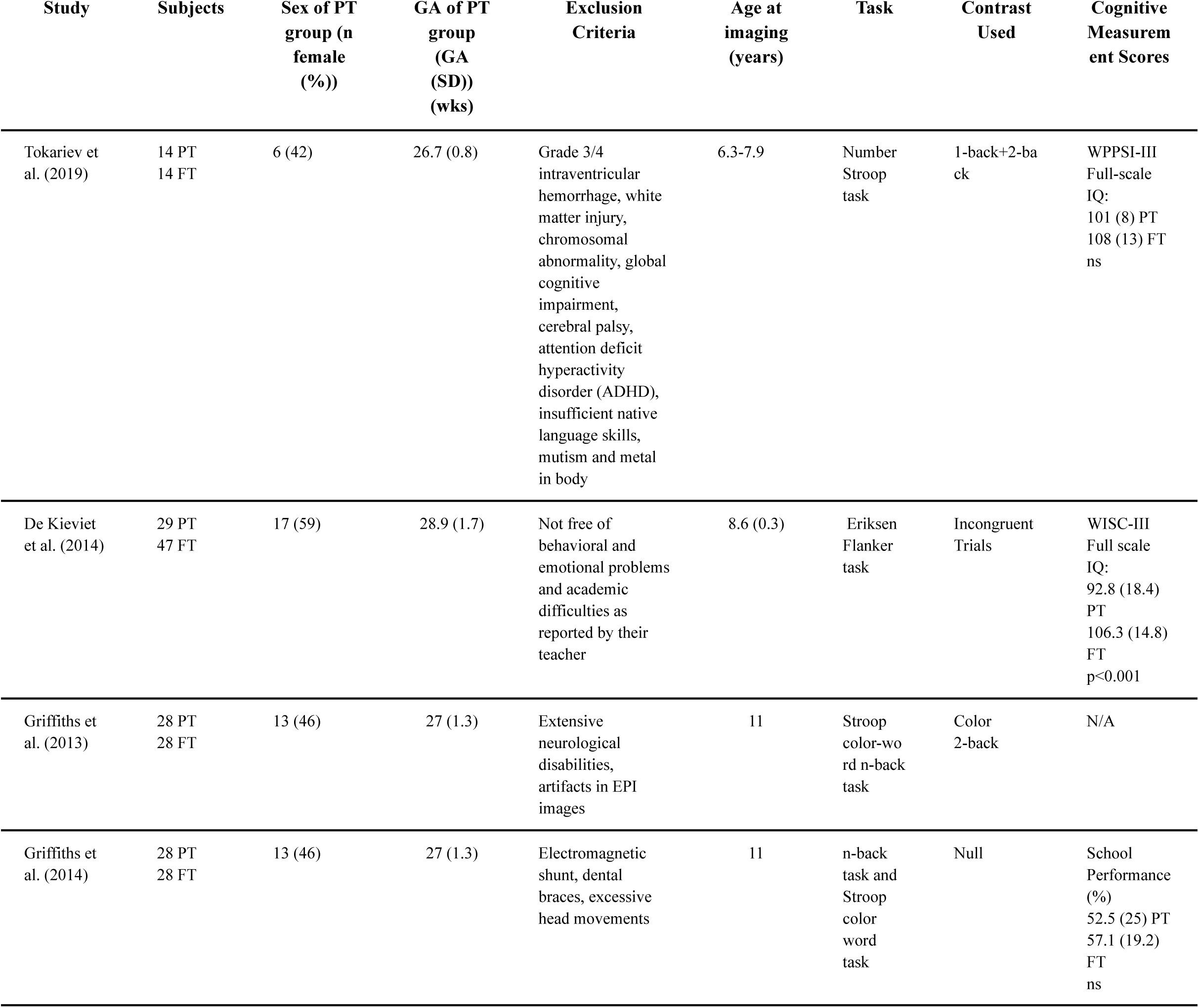
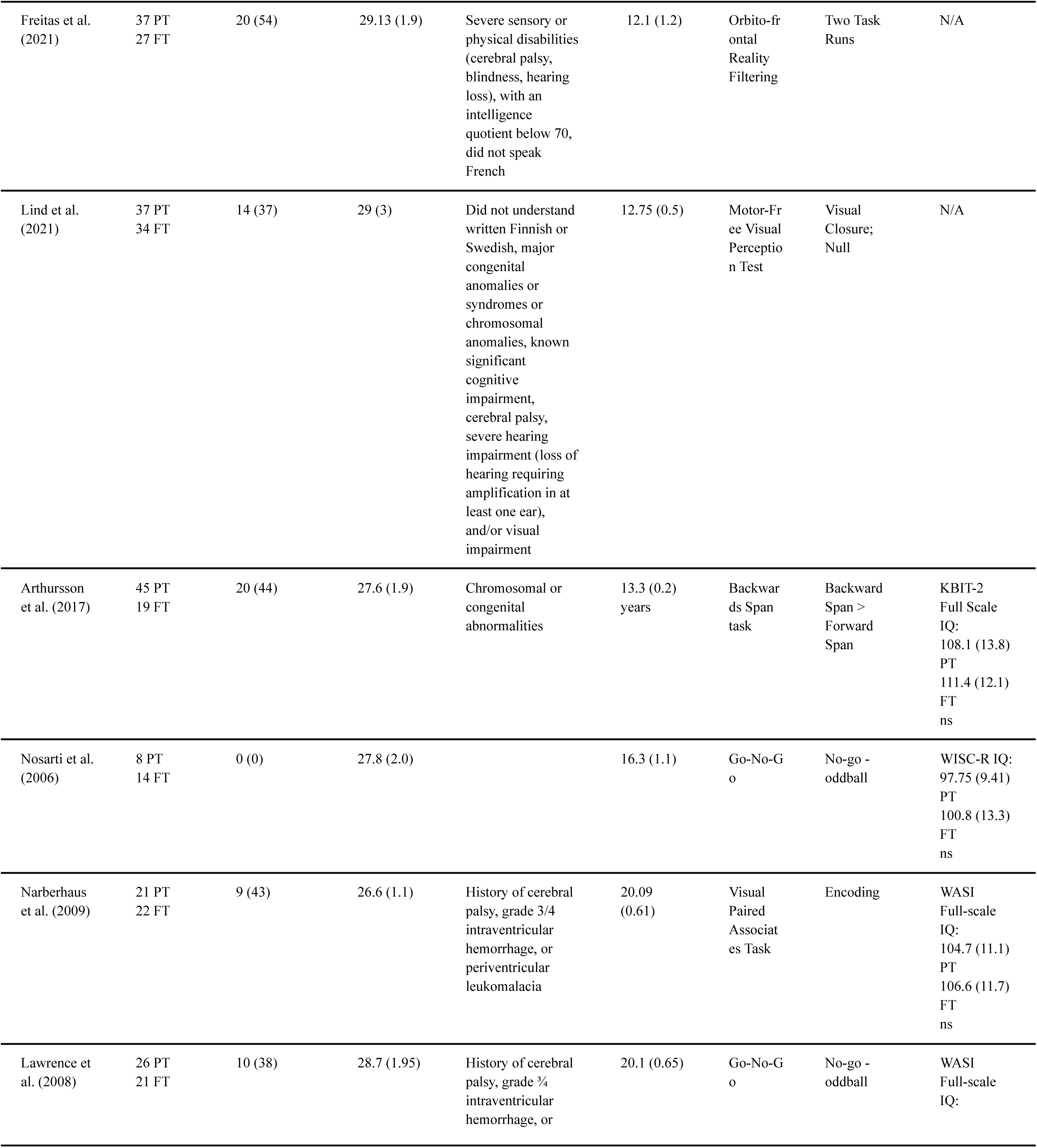
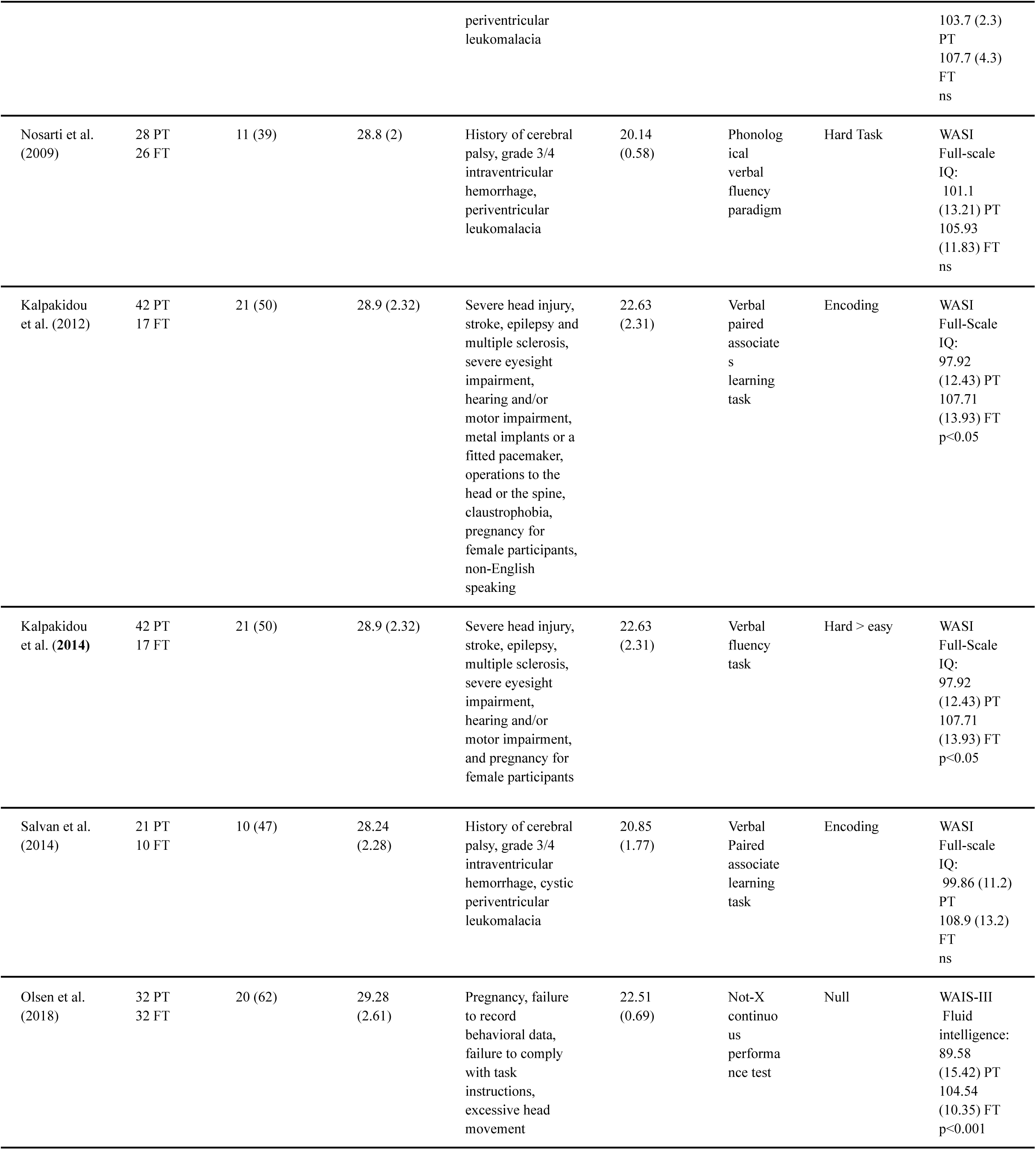
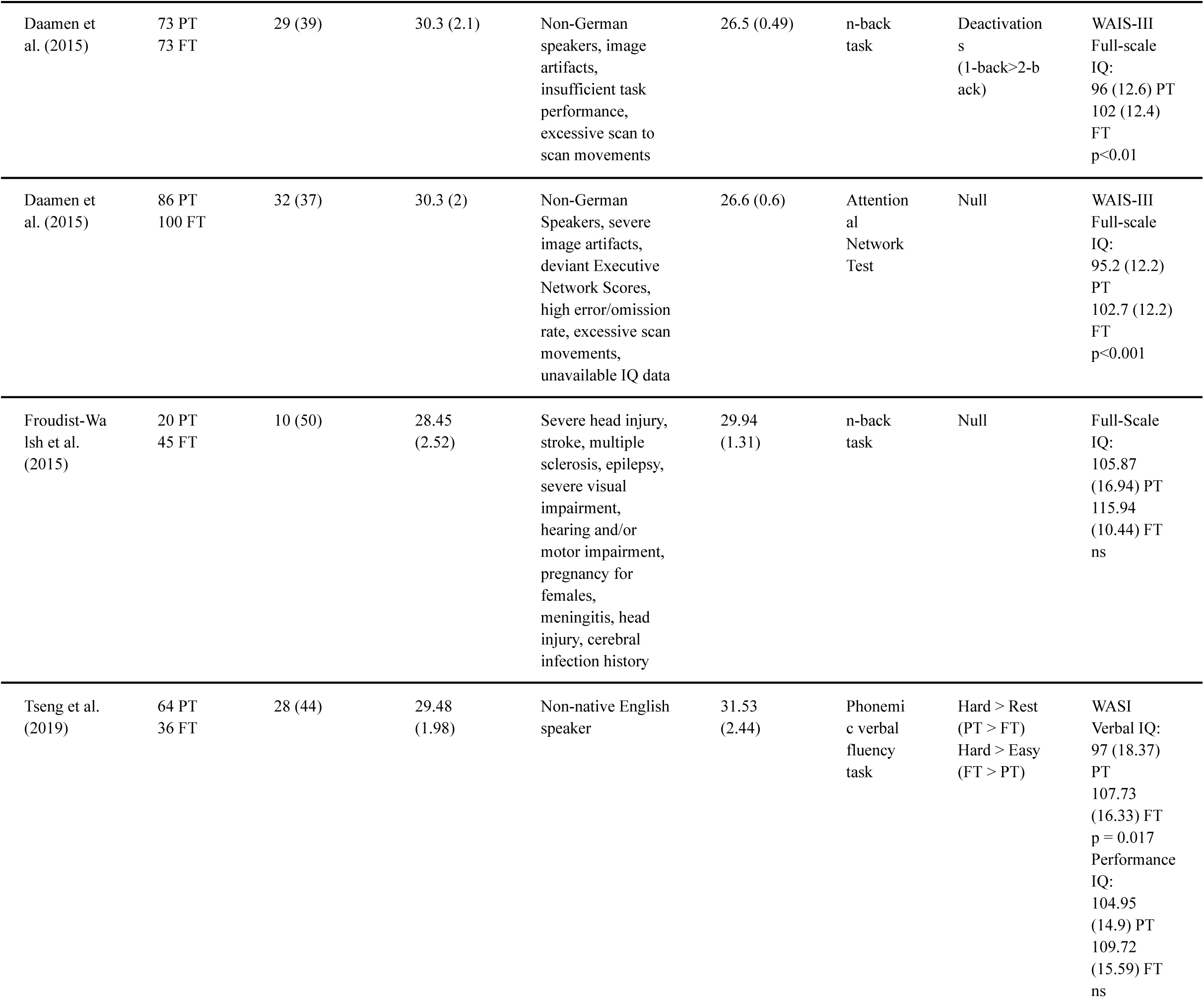
Studies included in Analysis.

### Data extraction

Two reviewers (AY, GS) independently assessed study quality and screened studies based on the above criteria. Both reviewers independently extracted reported voxels, and all disagreements were resolved by verbal consensus or using a tie breaker (CG) when necessary. Results were divided into two outcome measures: (1) voxels where PT participants displayed significantly lower activation compared to FT controls (FT>PT) and (2) voxels where PT participants displayed significantly greater activation than FT controls (PT>FT). If multiple studies used the same cohort of participants, only the voxels from the paper with the greatest number of reported foci were included in the analysis. The search was completed without excluding studies based on publication date or language used in the experimental tasks. All study review and data extraction were completed using Covidence (https://www.covidence.org/).

### Analysis

The MNI (x, y, z) coordinates for significant voxels served as the input for the meta-analysis. Each input included voxels taken from a single contrast in each study to avoid potential dependence across ALE maps (Turkeltaub et al., 2012). Included contrasts from each study are listed in Table 1. Any voxels reported in Talairach coordinates were converted to MNI coordinates prior to input. Cross study meta-analysis was completed using GingerALE version 2.3 (https://brainmap.org). GingerALE measures whether inter-experimental voxel convergence is higher than would be expected if results were independently distributed throughout the brain. GingerALE implements Turkeltaub’s method to reduce within-group effects and generates activation likelihood estimate (ALE) values for each voxel (Turkeltaub et al., 2002). These ALE values are used to generate the null distribution of each voxel. The generated p-values allow for the creation of a thresholded ALE map, and cluster analysis determines clusters of statistically significant overlap. A cluster-level familywise error rate (FWE), which accounts for potential Type I errors in the original data set, was set at <0.001. Each study underwent 1000 threshold permutations with threshold of p<0.05, as outlined in Müller et al., 2018.

ALE analyses were run for 6 different sets of results: 1) Adult results of hypoactivation in PT subjects compared to FT controls; 2) Adult results of hyperactivation in PT subjects compared to FT controls; 3) Child results of hypoactivation in PT subjects compared to FT controls; 4) Child results of hyperactivation in PT subjects compared to FT controls; 5) Contrast analysis between Adult and Child results; and 6) Conjunction analysis of Adult and Child results. Results were registered into MNI-template space and displayed using Mango (https://mangoviewer.com/).

## Results

### Characteristics of Included Studies

A total of 19 studies were included in this meta-analysis (see Table 1). Eight studies were in children ages 6-16, and eleven studies were in adults ages 20-35. The analysis included 198 PT children (two studies used the same subject population), 183 FT children, 413 PT adults (two adult studies used the same subject population), and 418 FT adults. 90 (49%) of child PT subjects and 180 (44%) of adult PT subjects were female. 16 of the 19 studies obtained behavioral cognitive measures from their study samples. 11 studies (4 child) measured full-scale IQ, 1 study measured school performance in children, 1 study measured fluid intelligence in adults, and 1 study measured verbal and performance IQ in adults. Differences between PT and FT groups were heterogeneous, with 10 of the 16 studies finding no significant differences in cognitive measures and the remaining six studies reported significantly lower scores in the PT group compared to the FT group. Across the studies with available full-scale IQ measurements, there was a significant difference (p<0.0001) between the group-weighted mean of full-scale IQ for PT subjects (*M*=99.3, *SD=*2.3) compared to FT subjects (*M*=106, *SD=*1.7). Details on the tasks utilized by each study, as well as the specific contrast utilized for data input for analysis are included in Table 1.

### ALE Results

#### Differences in Neural Engagement between PT and FT child subjects during Executive Functioning tasks

ALE analysis of voxels of significantly higher BOLD signal during EF tasks in child FT controls compared to PT subjects converged on one cluster, the right insula (see Fig. 1A). The cluster contained nine significant peaks (Table 2). ALE analysis of voxels representing regions of significantly higher BOLD activation in PT subjects compared to controls revealed one cluster of convergence in the left middle frontal gyrus (Fig. 1B). This cluster also contained nine significant peaks (Table 2).

**Figure 1.**
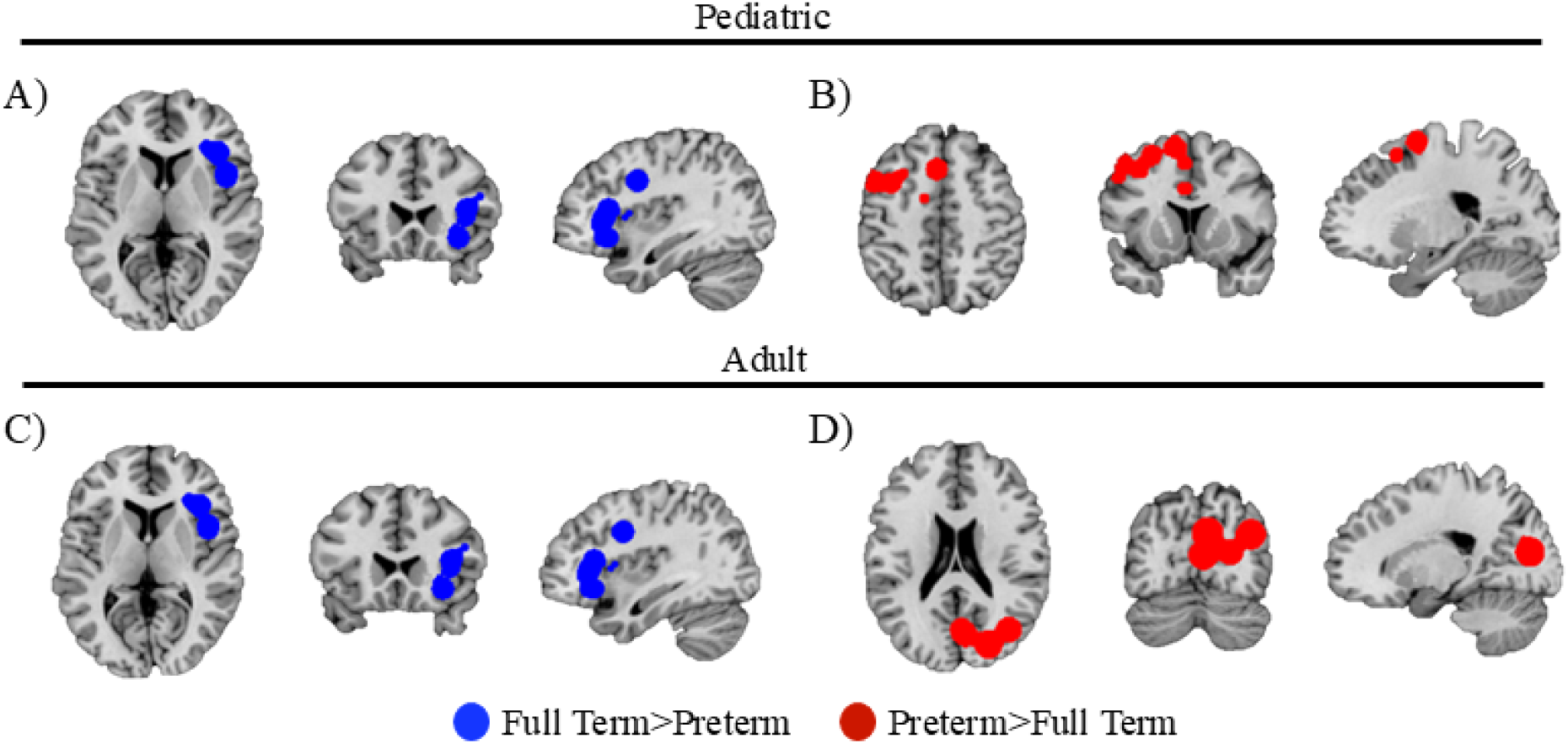
Comparison of convergence during EF tasks between PT and FT subjects. A) ALE analysis of voxels of greater BOLD signal in child FT subjects compared to PT subjects showed convergence in the right insula. B) Voxels of greater BOLD signal in child PT subjects compared to FT controls converged in the left middle frontal gyrus. ALE analysis of studies with adult subjects identified separate regions of convergence: C) voxels of greater BOLD signal in FT adults compared to PT adults show significant convergence in the right occipital gyrus, while D) voxels of greater BOLD signal in PT adults compared to FT adults converge in the right precuneus

**Table 2:** Results of ALE analysis.

| Cluster Label | Volume (mm³) | ALE value | P-value | Peak MNI coordinates |  |  |
| --- | --- | --- | --- | --- | --- | --- |
|  |  |  |  | *x* | *y* | *z* |
| Child, FT>PT |  |  |  |  |  |  |
| R Insula | 23024 | 0.009055 | <0.0001 | 32 | 26 | -10 |
|  |  | 0.008983 | <0.0001 | 38 | 26 | 4 |
|  |  | 0.008863 | 0.0001 | 22 | 42 | -8 |
|  |  | 0.008758 | 0.0002 | 36 | 2 | 30 |
|  |  | 0.008524 | 0.0002 | 46 | 8 | 8 |
|  |  | 0.008472 | 0.0002 | 40 | 22 | 10 |
|  |  | 0.00847 | 0.0002 | 46 | 12 | 4 |
|  |  | 0.007798 | 0.0007 | 30 | 30 | -2 |
|  |  | 0.005673 | 0.003404 | 48 | 12 | 20 |
| Child, PT>FT |  |  |  |  |  |  |
| L Middle Frontal Gyrus | 24016 | 0.009799 | <0.0001 | -36 | 8 | 44 |
|  |  | 0.009788 | <0.0001 | -20 | 0 | 64 |
|  |  | 0.00977 | <0.0001 | -10 | 12 | 60 |
|  |  | 0.009208 | <0.0001 | -48 | 8 | 48 |
|  |  | 0.0092 | <0.0001 | -12 | -2 | 54 |
|  |  | 0.009181 | 0.00018 | -4 | 18 | 48 |
|  |  | 0.00169 | 0.027515 | -6 | 24 | 27 |
|  |  | 0.008866 | 0.0002 | -28 | 14 | 54 |
|  |  | 0.008859 | 0.0003 | -50 | 18 | 40 |
| **Adult, FT>PT** |  |  |  |  |  |  |
| R Precuneus/Posterior Cingulate | 30592 |  |  |  |  |  |
|  |  | 0.010357 | <0.0001 | 16 | -62 | 22 |
|  |  | 0.009525 | <0.0001 | 14 | -64 | 12 |
|  |  | 0.008969 | 0.0002 | 12 | -66 | 38 |
|  |  | 0.008969 | 0.0002 | 6 | -44 | 26 |
|  |  | 0.008663 | 0.0003 | 10 | -42 | 26 |
|  |  | 0.001882 | 0.014413 | -1 | -43 | 28 |
| **Adult, PT>FT** |  |  |  |  |  |  |
| R Occipital gyrus/Cuneus | 27128 |  |  |  |  |  |
|  |  | 0.008632 | 0.0001 | 38 | -70 | 26 |
|  |  | 0.008632 | 0.0001 | 4 | -70 | 11 |
|  |  | 0.008346 | 0.0001 | 8 | -74 | 26 |
|  |  | 0.007969 | 0.0002 | 24 | -80 | 14 |

#### Differences in Neural Engagement between PT and FT Adults during Executive Functioning tasks

ALE analysis of studies reporting differences in neural engagement between FT and PT adults identified different clusters of significant overlap than the child analysis. Voxels of significantly higher BOLD signal in FT adults compared to PT adults converged in the right precuneus and posterior cingulate (Fig. 1C, Table 2). This cluster contained six significant peaks (Table 2). Voxels representing regions of increased BOLD activation during EF tasks in adult PT subjects revealed a significant cluster in the right occipital gyrus and cuneus (Fig. 1D). This cluster contained four significant peaks (Table 2).

#### Comparison between Adult and Child ALE Maps

Convergence maps between child and adult ALE maps for both PT>FT and FT>PT revealed no significant clusters, indicating that significant convergence in reported fMRI coordinates may only be relevant when considering the two age groups separately.

## Discussion

Our meta-analysis of studies investigating fMRI response during EF tasks revealed differences in the neural engagement during EF tasks in individuals born PT compared to those born FT. The fMRI data of decreased BOLD signal in PT children compared to FT children displayed significant convergence in the right insula, while fMRI data of increased BOLD signal in PT children had significant convergence in the left middle frontal gyrus. The right insula has been hypothesized to play a primary role in the salience network, integrating multisensory stimuli and detecting behaviorally-relevant information (Cai et al., 2014; Molnar-Szakacs & Uddin, 2022); decreased activity in salience network (SN) regions may cause PT children to have dampened ability to identify task-relevant sensory information. Hyperactivity in the left middle frontal gyrus supports the view of a dysregulated SN activity in PT children. The middle frontal gyrus is an important modulator of goal-directed behavior and identification of relevant sensory stimuli via fronto-striatal pathways (Grahn et al., 2008; Heitzeg et al., 2014). Abnormal functional connectivity between cortico-striatal pathways and SN nodes, such as the right insula, has been noted in various neuropsychiatric disorders affecting executive functioning, such as ADHD (Damiani et al., 2021; Silk et al., 2008) and OCD (Tomiyama et al., 2019). However, many of these studies report that inhibited executive functioning correlates with decreased activity in fronto-striatal pathways (Durston et al., 2003) and greater fronto-striatal activity correlates with improved behavioral responses in typically developing populations (Durston et al., 2002; Rubia et al., 2006). Therefore, hyperactive fronto-striatal activity seen during EF tasks in PT children’s pathways may be a compensatory response to decreased SN activity.

Importantly, the left middle frontal gyrus has consistently been identified as a general executive functioning region, as it is consistently noted as a significant region of activation in conjunction analyses across EF subdomain tasks (Collette et al., 2005; Saylik et al., 2022; Sylvester et al., 2003). The regions identified in our child results are related to broader EF capabilities. These results are consistent with the hypothesis that PT birth may impact broader EF networks as opposed to networks supporting specific EF subdomains. Of particular relevance in childhood seems to be the dysregulation between fronto-striatal networks and the salience network.

Our adult ALE results identified unique significant clusters differing between PT and FT subjects. PT adult compared to FT adult fMRI data of decreased BOLD signal had significant convergence in the right precuneus and posterior cingulate. The clusters in the precuneus and posterior cingulate–are key components of the default mode network (DMN) (Raichle, 2015; Shulman et al., 1997). Greater suppression of the DMN, which typically shows decreased activity during cognitively demanding tasks, is related to improved performance (Whitfield-Gabrieli & Ford, 2012). However, there is evidence that in younger adults, greater hypoactivation of the DMN correlates with worse performance (Grahn et al., 2008). This decreased DMN activity may reflect greater effort exertion during the EF tasks (Weber et al., 2022), indicating the tasks may be more difficult for PT subjects. Importantly, DMN deactivations are seen across EF tasks, regardless of the subdomain the tasks target (Reineberg et al., 2022), implying that changes in DMN recruitment would impact broad EF skills as opposed to individual subdomains.

Our adult ALE results also revealed regions where fMRI data of increased BOLD signal in PT adults compared to FT adults was observed. Specifically, significant convergence in the right occipital gyrus and cuneus, areas primarily involved in early visual processing (Vanni et al., 2001; Zeng et al., 2020). These regions are not typically associated with core executive function (EF) networks but are functionally linked to the visual system and may show co-activation with attention networks like the dorsal attention network (DAN) during visually demanding tasks (Corbetta & Shulman, 2002; Vossel et al., 2014). Elevated activity in these areas may reflect compensatory visual processing strategies or greater reliance on bottom-up visual input to support task performance. This shift in activation may reflect compensatory recruitment of perceptual resources in PT adults, potentially due to inefficiencies in higher-order control networks. Notably, similar visual region recruitment has been observed in other populations during increased task difficulty or when top-down systems are underdeveloped or impaired (Menon, 2011), suggesting that visual hyperactivation in PT adults may reflect broader alterations in cognitive strategy or neural resource allocation (Narberhaus et al., 2009; Froudist-Walsh et al., 2015; Tseng et al., 2016).

Overall, our meta-analysis suggests that significant differences in neural engagement during EF tasks exist in both children and adults born PT compared to FT controls. Despite negative contrast analyses, there also seem to be important differences between the regions implicated in PT children and adults. While PT children show hyperactivity in fronto-striatal regions, brain regions implicated in PT adults suggest changes to default mode network activity. The DMN is not fully matured in children and early adolescents (de Bie et al., 2011; Mak et al., 2017; Thomason et al., 2008), which may explain why we see no significant differences emerge in DMN regions in PT children. PT adults also show increased activation in posterior cortical regions, including early visual areas such as the right occipital gyrus and cuneus (Vanni et al., 2001; Zeng et al., 2020), which may reflect broader differences in neural strategy or task engagement that emerge with age (Narberhaus et al., 2009; Froudist-Walsh et al., 2015; Tseng et al., 2016). However, this may also reflect the increased proportion of visually-associated tasks represented in our adult studies compared to the child studies. It is important to note that EF task modalities were unevenly represented across studies, which may bias results toward more frequently studied tasks. While this limits subdomain-specific conclusions, we observed consistent involvement of core EF-related regions, which may reflect broader executive processing differences. Both child and adult results are consistent with the possibility that activity in regions subserving shared EF components are impacted by PT birth.

## Limitations and Future Directions

While the current study provides deeper insight into the general differences in the brain networks subserving EF in individuals born prematurely, there are limitations to the current body of literature. Firstly, all the studies included in our meta-analysis focus on a limited subset of the preterm population. Every study exclusively focused on very preterm (<32 weeks GA) individuals, excluding moderately and late preterm individuals (32-37 weeks GA), despite that this group accounts for the majority of preterm births and also displays risk for deficits in cognitive functions (Baron et al., 2012; Hodel et al., 2017; Talge et al., 2010). Future studies should aim to expand GA criteria or even exclusively focus on these late preterm individuals to better characterize EF outcomes across the heterogeneous population of individuals born prematurely. Additionally, the current sample is not matched on general IQ measurements. As a result, it is not possible to say whether differences in EF processing in PT individuals are compensatory or are potential targets for intervention. Future work should expand on whether the differences in neural activity covered in this work relates to behavior during EF tasks. Finally, the range of different EF tasks included in the current study is not complete due to a lack of studies including cognitive flexibility or shifting. While we do not believe the focus on working memory, inhibition, and broad EF tasks diminishes the impact of our findings, future work would benefit from including a more expansive battery of EF tasks covering each subdomain.

## Conclusions

Overall, this meta-analysis has identified areas of differential activation during EF tasks in individuals born PT both in childhood and adulthood. Our results suggest that salience network regions are affected in both children and adults, identifying a locus of persistent differential executive processing in individuals born preterm. However, fronto-striatal networks may be uniquely affected in children, while default mode network regions were identified in adults. These results provide evidence that PT birth impacts EF through domain-wide alterations in brain function. Further work should aim to build upon these findings through fMRI studies targeting specific EF subdomains.

